# The prevalence of *Fusobacterium nucleatum* subspecies in the oral cavity stratifies by local health status

**DOI:** 10.1101/2023.10.25.563997

**Authors:** Madeline Krieger, Yasser M. AbdelRahman, Dongseok Choi, Elizabeth A. Palmer, Anna Yoo, Sean McGuire, Jens Kreth, Justin Merritt

## Abstract

The ubiquitous inflammophilic pathobiont *Fusobacterium nucleatum* is widely recognized for its strong association with a variety of human dysbiotic diseases such as periodontitis and oral/extraoral abscesses, as well as multiple types of cancer*. F. nucleatum* is currently subdivided into four subspecies: *F. nucleatum* subspecies *nucleatum* (*Fn. nucleatum*)*, animalis* (*Fn. animalis), polymorphum* (*Fn. polymorphum*), and *vincentii/fusiforme* (*Fn. vincentii*). Although these subspecies have been historically considered as functionally interchangeable in the oral cavity, direct clinical evidence is largely lacking for this assertion. Consequently, we assembled a collection of oral clinical specimens to determine whether *F. nucleatum* subspecies prevalence in the oral cavity stratifies by local oral health status. Patient-matched clinical specimens of both disease-free dental plaque and odontogenic abscess were analyzed with newly developed culture-dependent and culture-independent approaches using 44 and 60 oral biofilm/tooth abscess paired specimens, respectively. Most oral cavities were found to simultaneously harbor multiple *F. nucleatum* subspecies, with a greater diversity present within dental plaque compared to abscesses. In dental plaque, *Fn. polymorphum* is clearly the dominant organism, but this changes dramatically within odontogenic abscesses where *Fn. animalis* is heavily favored over all other fusobacteria. Surprisingly, the most commonly studied *F. nucleatum* subspecies, *Fn. nucleatum,* is only a minor constituent in the oral cavity. To gain further insights into the genetic basis for these phenotypes, we subsequently performed pangenome, phylogenetic, and functional enrichment analyses of oral fusobacterial genomes using the Anvi’o platform, which revealed significant genotypic distinctions among *F. nucleatum* subspecies. Accordingly, our results strongly support a taxonomic reassignment of each *F. nucleatum* subspecies into distinct *Fusobacterium* species. Of these, *Fn. animalis* should be considered as the most clinically relevant at sites of active inflammation, despite being among the least characterized oral fusobacteria.

## Introduction

*Fusobacterium nucleatum* is a ubiquitous member of the human oral microbiome, where its distinctive filamentous structure and plethora of interspecies interactions allow it to act as a bridging organism to facilitate the integration of numerous late colonizing microorganisms into the oral biofilm (1, 2). *F. nucleatum* has also long been recognized for its prominent role in a variety of oral diseases, especially odontogenic abscesses and periodontitis (3). Similarly, *F. nucleatum* has garnered widespread attention due to its prominent association with multiple extraoral conditions, including preterm birth, head and neck cancers, colorectal cancer (CRC), and others (4–11). *F. nucleatum* has been shown to target specific innate immune signaling pathways and modulate the autophagic response, leading to enhanced chemoresistance in CRC (12). Despite its association with a broad array of oral and extraoral diseases, substantial knowledge gaps still remain in our mechanistic understanding of *F. nucleatum* pathobiology.

Up to five subspecies of *F. nucleatum* have been previously described: *F. nucleatum* subspecies *nucleatum* (*Fn. nucleatum*)*, F. nucleatum* subsp. *animalis* (*Fn. animalis), F. nucleatum* subsp. *polymorphum* (*Fn. polymorphum*), and *F. nucleatum* subsp*. vincentii/fusiforme*, which are both genetically identical and are now considered as a single subspecies (henceforth referred to as *Fn. vincentii*) (13–16). Isolates from all four of these subspecies, as well as the closely related species *F. periodonticum*, have been cultured from the oral cavity and are typical members of the human microbiota. Furthermore, identical strains of *F. nucleatum* have been isolated from both colorectal tumors and the oral cavities of CRC patients, implicating the oral cavity as the primary source of *F. nucleatum* strains seeding tumor sites (17, 18).

To date, *F. nucleatum* subspecies have been treated as largely interchangeable in laboratory investigations and clinical studies. This may be partially attributable to the fact that subspecies classification of *F. nucleatum* isolates is often challenging using typical 16S sequencing approaches, due to the extensive sequence identity shared by their 16S rRNA genes. Consequently, there are a variety of fundamental unanswered questions regarding *F. nucleatum* prevalence in health and disease. For example, it remains unclear which, if any, subspecies predominate within the typical human microbiota, nor is it known whether health and/or disease association stratifies with *F. nucleatum* subspecies. Given the recent evidence implicating *Fn. animalis* as the predominant subspecies associated with CRC tumors (4, 10, 19) as well as its likely origination in the oral cavity (17, 20), these questions are particularly timely.

To address these knowledge gaps, we developed new culture-dependent and culture-independent genotyping approaches to identify and/or quantify *F. nucleatum* subspecies directly from clinical specimens. Using both approaches, we investigated the prevalence of *F. nucleatum* subspecies found in patient-matched clinical specimens of health and disease. Our results reveal striking biases in *F. nucleatum* subspecies composition among the paired dental plaque and abscess clinical specimens, suggesting fundamental differences likely exist in the ecological niches preferred by these organisms as well as their potential pathogenicity. Furthermore, whole-genome phylogenetic and pangenome comparisons of *F. nucleatum* subspecies identified significant divergences between clades currently classified as subspecies. In contrast to the current view of this organism, our results provide multiple lines of evidence indicating that the four *F. nucleatum* subspecies are not relatively synonymous organisms, rather they likely comprise unique *Fusobacterium* species having distinct roles in human health and disease.

## Results

### 1. Culture-dependent analysis of *F. nucleatum* subspecies within the patient cohort

While there is a plethora of studies demonstrating the ubiquity of *F. nucleatum* among human oral microbiomes (1), the diversity of oral fusobacteria within individual oral cavities has remained unclear, nor is it known whether particular *F. nucleatum* subspecies predominate at sites of health and disease. These questions are of interest because it is often assumed that the different *F. nucleatum* subspecies exist somewhat interchangeably, as might be expected from closely related subspecies. To address these questions as well as simplify *F. nucleatum* subspecies identifications of clinical isolates, our first aim was to develop a rapid PCR-based identification approach that would abrogate the requirement for DNA sequencing. To this end, we developed a PCR primer set that quickly and unambiguously discriminates between the four *F. nucleatum* subspecies as well as *F. periodonticum* due to the production of uniquely sized subspecies-specific PCR amplicons (**Figure 1A**). The specificity of these primers was tested *in silico* using 58 *F. nucleatum* genomes downloaded from NCBI (**Table S1 and S2**). Prior to analyzing our clinical specimen collection, we first confirmed that this approach could reliably detect a single *Fusobacterium* colony when scraped from an agar plate together with >500 CFU of other species. Since this method performed as expected and further allowed us to rapidly identify the species/subspecies of individual isolates, we next employed this strategy to examine the composition of fusobacteria present within the oral patient specimens (**Figure 1B**). To stratify the results based upon oral health status, we compared patient-matched clinical specimens concurrently sampled from dental plaque collected from caries-free teeth vs. odontogenic abscess material collected from extracted abscessed teeth. The patient specimens were collected from a cohort of 44 pediatric dental patients previously scheduled for tooth extraction due to one or more tooth abscesses. Abscess clinical specimens were sampled by swabbing purulence from the furcation or root tip of a tooth following its extraction. Odontogenic abscess samples were specifically chosen for analysis due to their physical sequestration away from the oral biofilm, thus minimizing the opportunity for cross contamination of samples during their collection. We plated paired patient plaque and abscess specimens directly onto selective agar plates, extracted genomic DNA (gDNA) from the pooled collection of colonies scraped from each plate, and then performed PCR using the primers listed in Table 1. When using a qualitative assessment of presence/absence counts derived from these pooled colonies, we were somewhat surprised to find that most patient specimens simultaneously harbored multiple types of fusobacteria. The PCR results suggested *F. periodonticum* is primarily found within the dental plaque specimens, but other biases were not apparent (**Figure 2, Table S3**). However, we also noted that this approach is of limited utility to reliably discriminate overall abundances in samples containing multiple fusobacteria. Furthermore, while collecting individual fusobacterial clinical isolates from the selective plates, we unexpectedly observed what appeared to be a highly biased distribution of fusobacteria in the patient-matched oral plaque and abscess specimens. Individual *F. periodonticum* and *Fn*. *polymorphum* isolates were identified far more frequently on the plates receiving dental plaque specimens, whereas *Fn*. *animalis* was the principal organism isolated from selective plates spread with odontogenic abscess material. Thus, our observational results suggested that the distribution of *Fn* subspecies within plaque and abscess specimens could be far more biased than the pooled qualitative sample analysis suggested (**Figure 2**). For this reason, we were interested to develop a quantitative culture-independent analysis as a more direct comparison of fusobacterial abundance in patient plaque and abscess clinical specimens.

**Figure 1.**
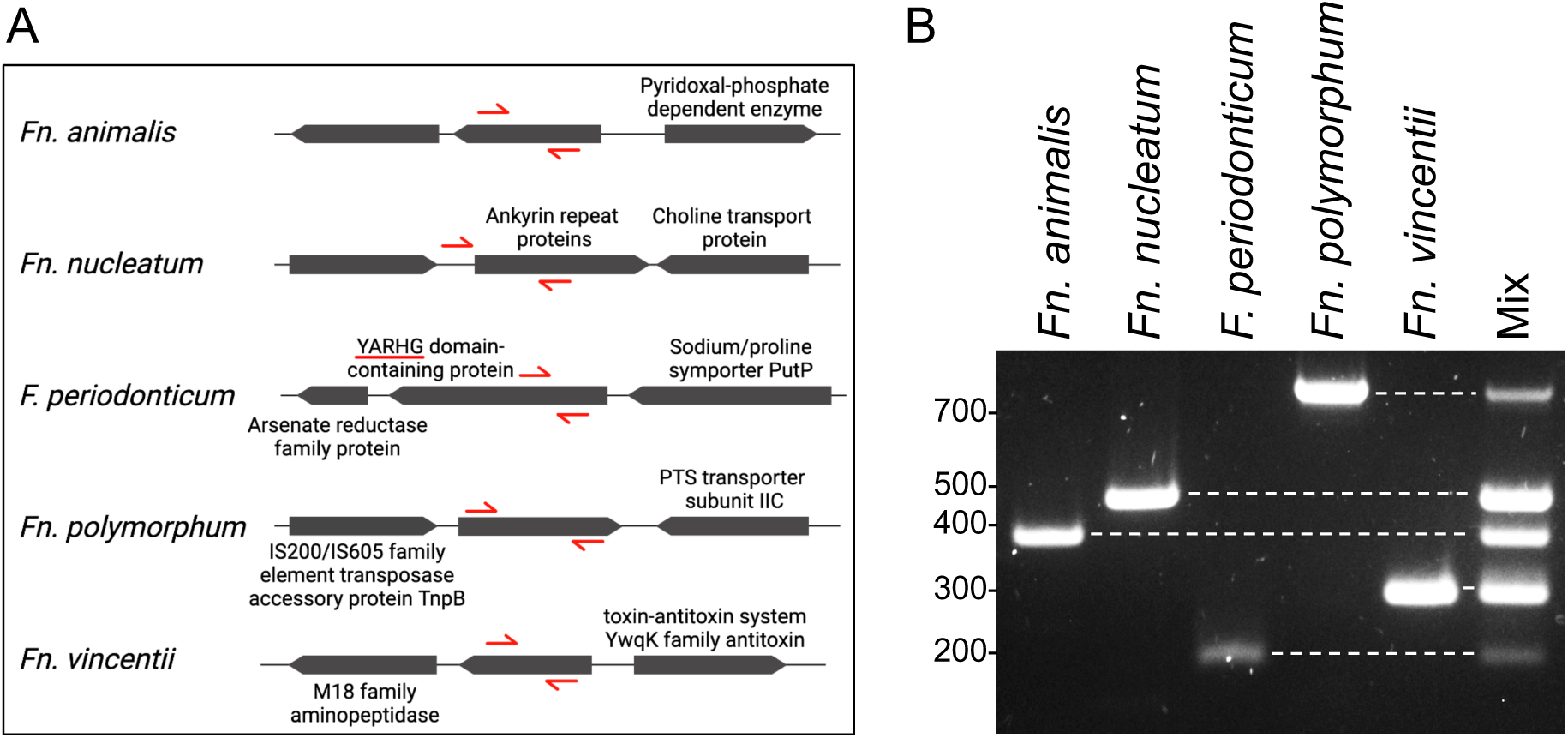
Subspecies determination using PCR primers. A) Location of subspecies-/species-specific primer sets in each *Fusobacterium* genome. Genes are labeled with their annotated function if available, while hypothetical proteins are unlabeled. Forward and reverse primer locations are illustrated with red arrows. Genes and primers are not drawn to scale. B) Representative example of subspecies/species primer specificity. All reactions were performed using a mixture of gDNA from the four *F. nucleatum* subspecies and *F. periodonticum*. A DNA ladder is shown on the left.

**Figure 2.**
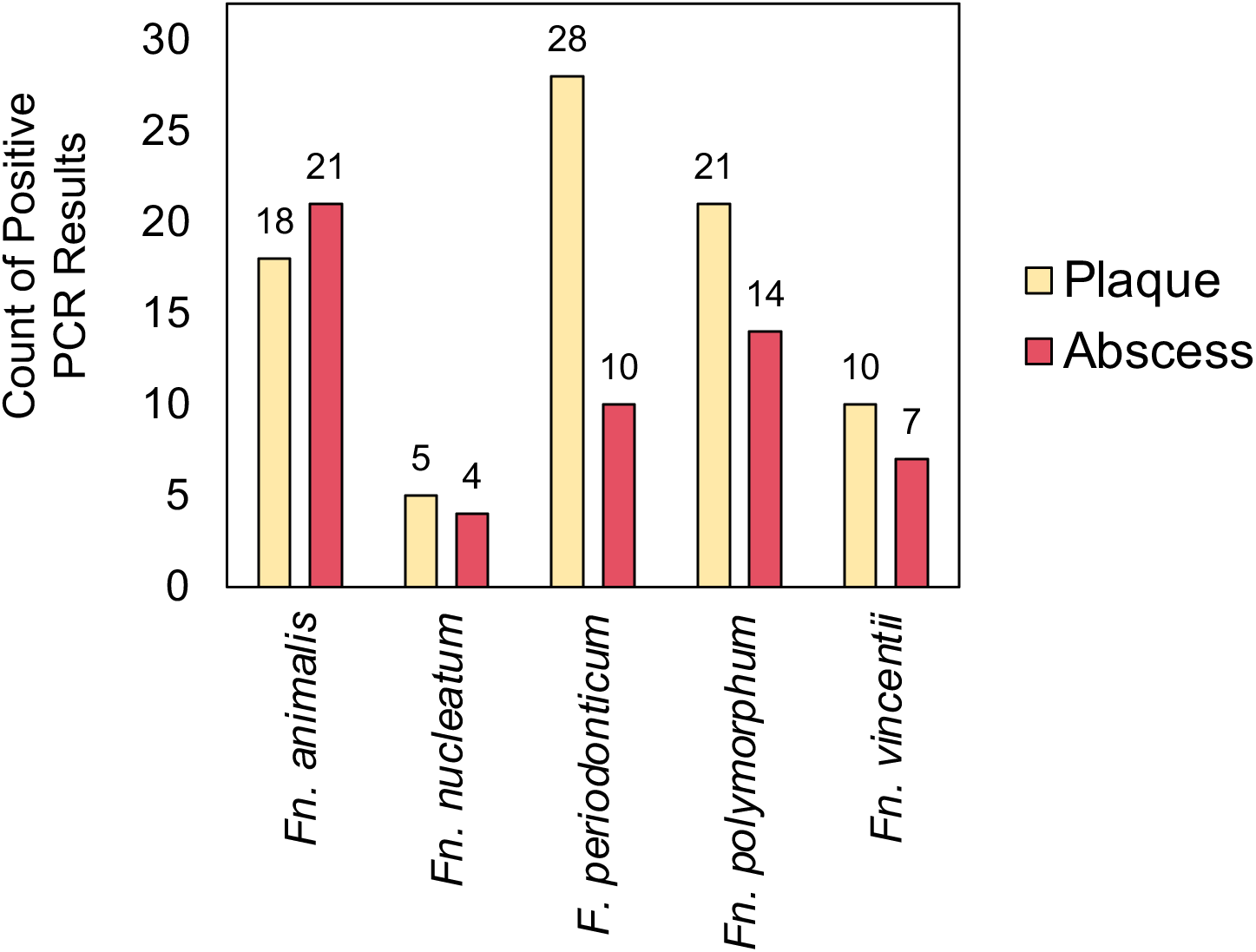
Detection of *Fusobacterium* species/subspecies in plaque and abscess samples using subspecies/species specific PCR primers. A total of 44 patient-matched plaque and abscess specimens were spread onto enrichment plates before pooling the CFU from each plate to survey for the presence or absence of each species/subspecies.

**Table 1.**
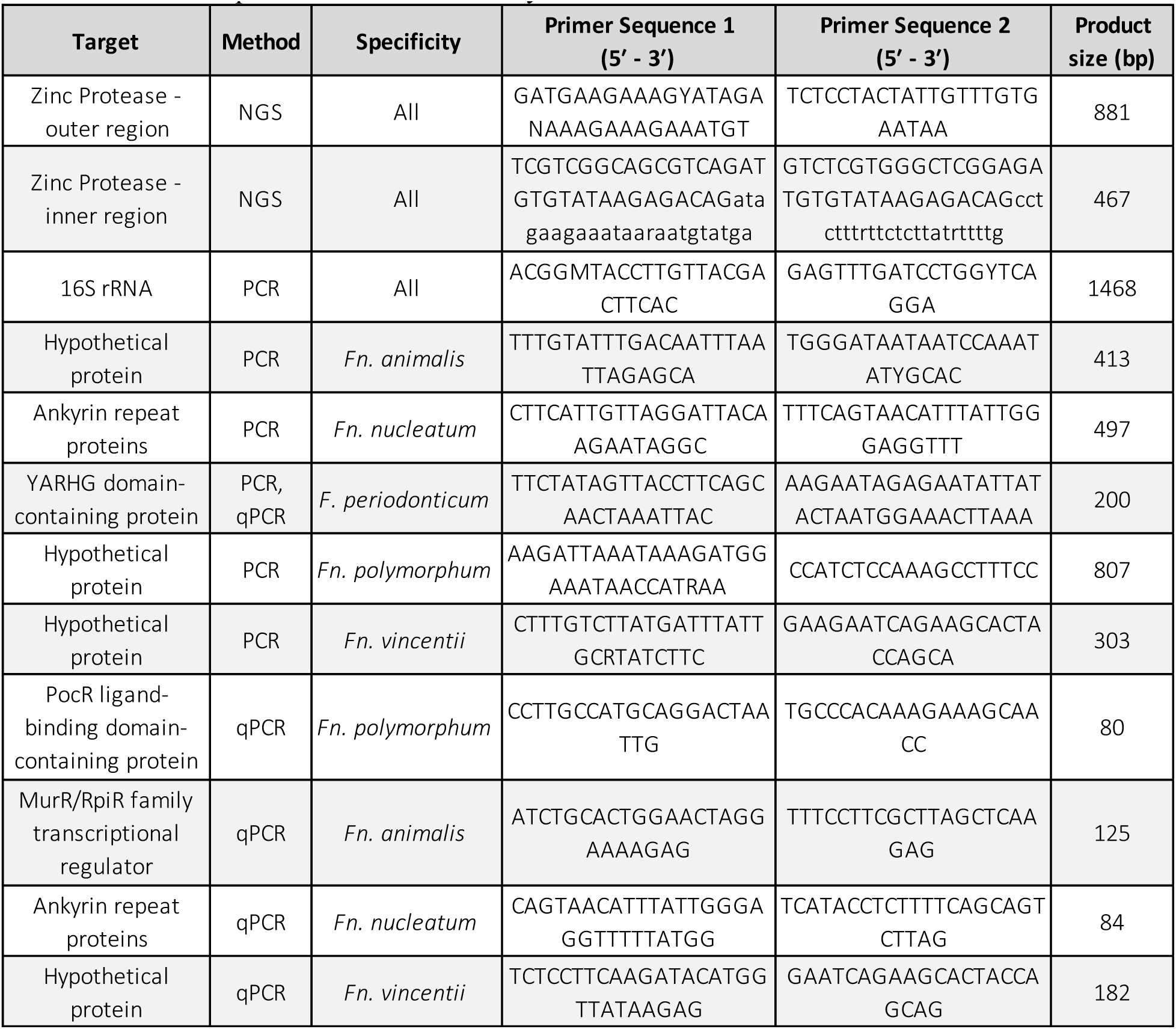
Primer sequences used in this study.

### 2. A hypervariable region of the fusobacterial *znpA* gene facilitates *F. nucleatum* subspecies-level classifications

Most quantitative NGS-based bacterial community analyses rely upon alignments of the 16S rRNA gene hypervariable V3V4 region. However, this approach can be highly unreliable when classifying closely related organisms, such as *F. nucleatum* subspecies **(Table 2, Figure S1)**. Previous work demonstrated that orthologs of the highly conserved fusobacterial gene encoding the zinc protease (henceforth referred to as *znpA*) contains sufficient sequence diversity to support accurate and reliable subspecies typing of *F. nucleatum* clinical isolates (21). This prior study utilized the entire *znpA* sequence (878 bp), which is impractical to employ for typical short-read Illumina-based NGS approaches. Therefore, we identified a 467 bp hypervariable segment of *znpA* flanked by highly conserved primer-binding sites (**Table S3**) as a potential alternative target for Illumina-based NGS of *F. nucleatum* subspecies. As shown in **Table 2** and **Figure S1**, sequence alignments identified >10-fold more single nucleotide polymorphisms within the identified *znpA* hypervariable region compared to the fusobacterial 16S rRNA V3V4 region. As a final confirmation, we compared the *znpA* hypervariable sequences from 58 sequenced *F. nucleatum* strains listed in the NCBI database (**Table S1**) and confirmed that strains could be accurately classified to the species and subspecies levels using the resulting *znpA* PCR amplicon (**Table S4**). Consequently, we next sought to sequence this same *znpA* hypervariable region directly from an expanded cohort of clinical specimens to quantify the proportions of oral fusobacteria contained within.

**Table 2.** Comparison of nucleotide variability between *Fusobacterium* species/subspecies within the zinc protease (*znpA*) gene sequence and the 16S V3V4 region.

|  | Segment Length | Number of SNPs | Average number of nucleotides between SNPs |
| --- | --- | --- | --- |
| **Zinc Protease** | 466 | 104 | 3.5 |
| **16S V3v4** | 442 | 9 | 43.3 |

### 3. Culture-independent quantitation of oral fusobacteria within patient-matched dental plaque and odontogenic abscess specimens

Initially, the limited quantities of patient material present within individual dental specimens were complicated by the inherently low abundance of bacterial DNA present within the samples. Therefore, we developed a nested PCR approach using two rounds of PCR amplification to generate sufficient DNA for NGS sequencing (**Table 1**). Before analyzing the resulting Illumina sequence data, we first imposed a detection threshold of at least 10,000 unique *znpA* reads among the individual samples to eliminate the low confidence samples from the dataset, resulting in 60 plaque and abscess specimen pairs meeting our inclusion criteria. From these data, it was clear that both plaque and abscesses typically harbor multiple types of fusobacteria, with most exhibiting highly biased distributions between the two sample sites (**Figure 3A**, **Table 3**). *Fn. polymorphum* and *Fn. animalis* comprised the most abundant oral fusobacteria overall, but they also exhibited distinct inversely proportional abundances between the plaque and abscess specimens (**Figure 3B**). *Fn. polymorphum* clearly predominates within plaque samples, comprising ∼50% of the total collection of *znpA* sequence reads, but this level drops precipitously in abscesses (**Figure 3B**). In contrast, ∼50% of all *znpA* sequence reads in the abscesses could be attributed to *Fn. animalis*, whereas a far smaller proportion was found within plaque specimens (**Figure 3B**). These biases were even more pronounced when compared on a per patient individual basis, rather than as a pooled collection (**Figure 3A**). *Fn*. *vincentii* was the only organism exhibiting no obvious biases, as it was detected at nearly identical rates in both plaque and abscess samples, contributing ∼20% of all sequence reads in each. *F. periodonticum* and *Fn*. *nucleatum* were the least abundant fusobacteria detected overall in both plaque and abscess samples. However, in the minority of specimens in which they were detected, both organisms also exhibited highly biased distributions. *F. periodonticum* is far more abundant within dental plaque, whereas *Fn*. *nucleatum* prefers the abscess environment (**Figure 3B**). Overall, these results indicate that most oral fusobacteria are highly unlikely to be randomly distributed amongst sites of health and disease.

**Figure 3.**
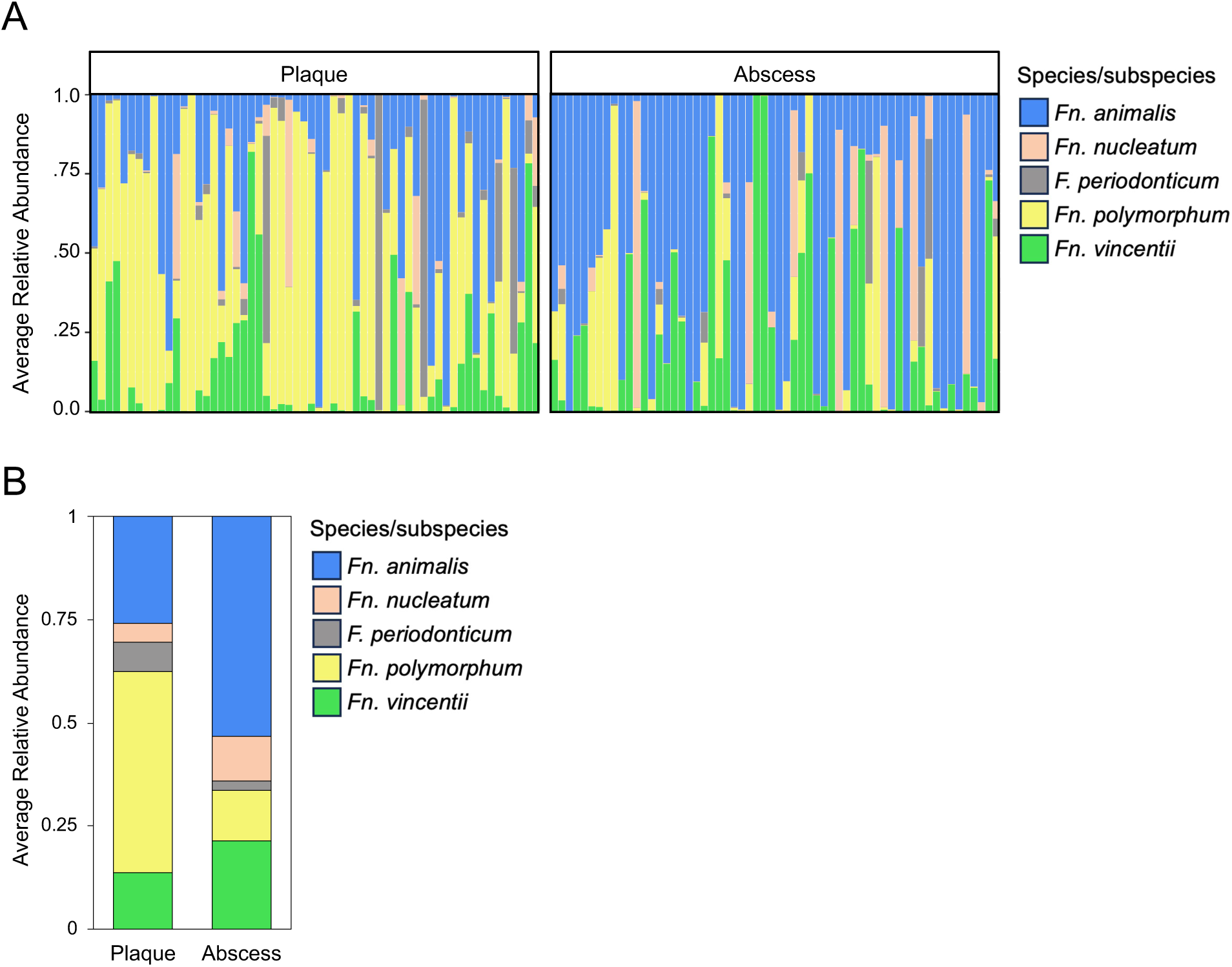
NGS quantitation of *Fusobacterium* species/subspecies in paired dental plaque and abscess samples. A) Relative abundances of 60 paired plaque and abscess specimens indicate distinct biases favoring *Fn. polymorphum* in plaque and *Fn*. *animalis* in odontogenic abscesses. Each column represents an individual patient specimen, with fusobacterial abundances presented proportionally. The plaque and abscess results are arranged in identical order, and are color coded according to *Fusobacterium* species/subspecies. B) Total relative abundances of each *Fusobacterium* species/subspecies among the entire collection of 60 paired dental plaque and abscess specimens.

**Table 3.**
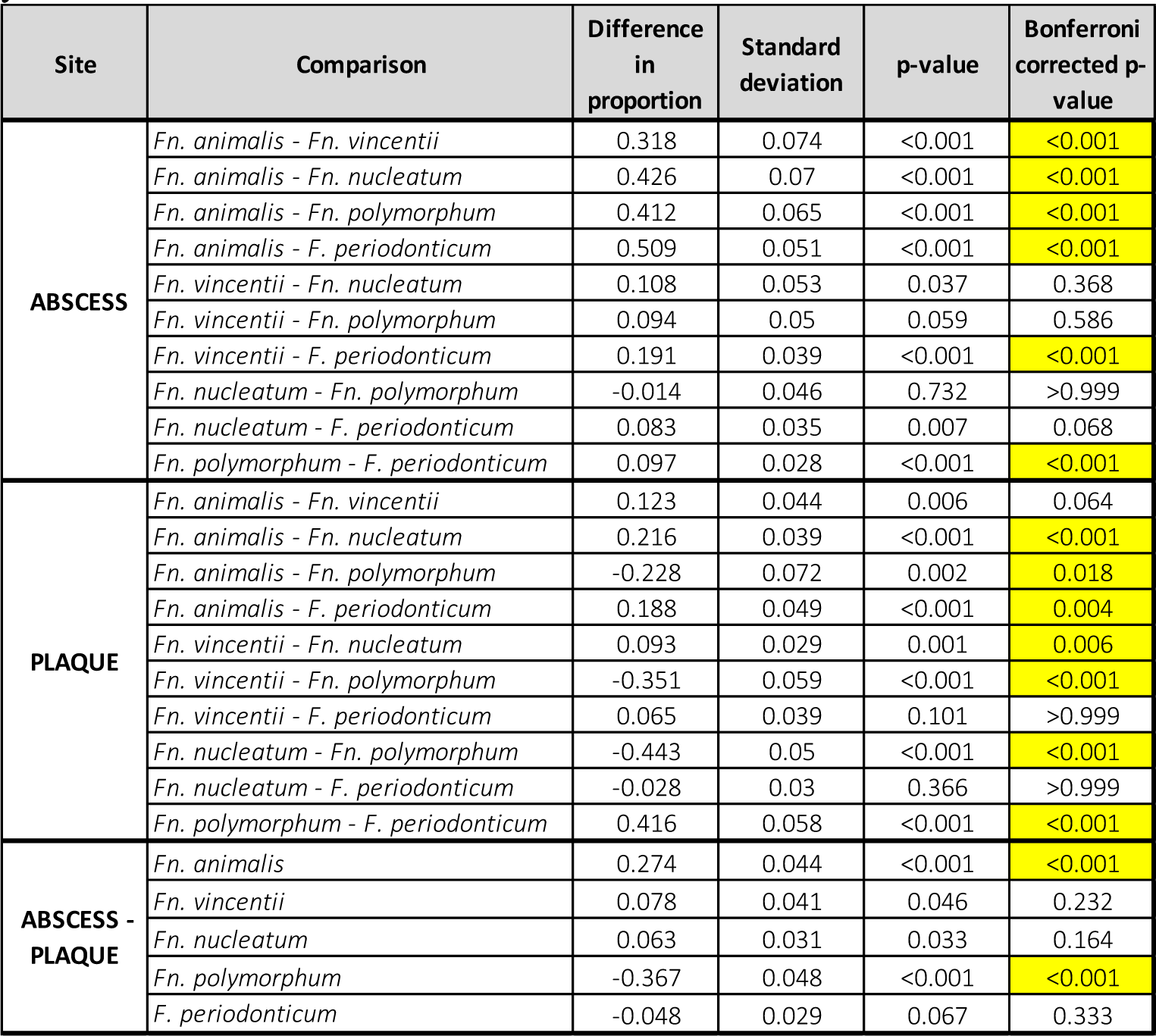
Comparison of fusobacterial proportions within clinical specimens using the NGS identification approach. Significant Bonferroni corrected *p*-values (p<.05) are highlighted in yellow.

### 4. Comparison of fusobacterial diversity in dental plaque and odontogenic abscesses

To date, the number of oral *Fusobacterium* species/subspecies normally harbored within an individual oral cavity has remained unclear. To answer this fundament question, we utilized the NGS data to determine the average co-occurrence of fusobacterial species/subspecies within dental plaque and abscess samples. We set a threshold of 1% total fusobacterial species/subspecies abundance values within each sample as being indicative of stable colonization in the host as part of the resident microbiota. Using this criterion, >87% of the cohort were confirmed to harbor multiple distinct *Fusobacterium* species/subspecies within dental plaque (**Figure 4**), with >33% simultaneously hosting 4 – 5 different types. Most abscess specimens also contained a mixture of multiple species/subspecies, but the diversity was noticeably lower overall, with >60% of the specimens containing only 1 or 2 distinct fusobacteria (**Figure 4**).

**Figure 4.**
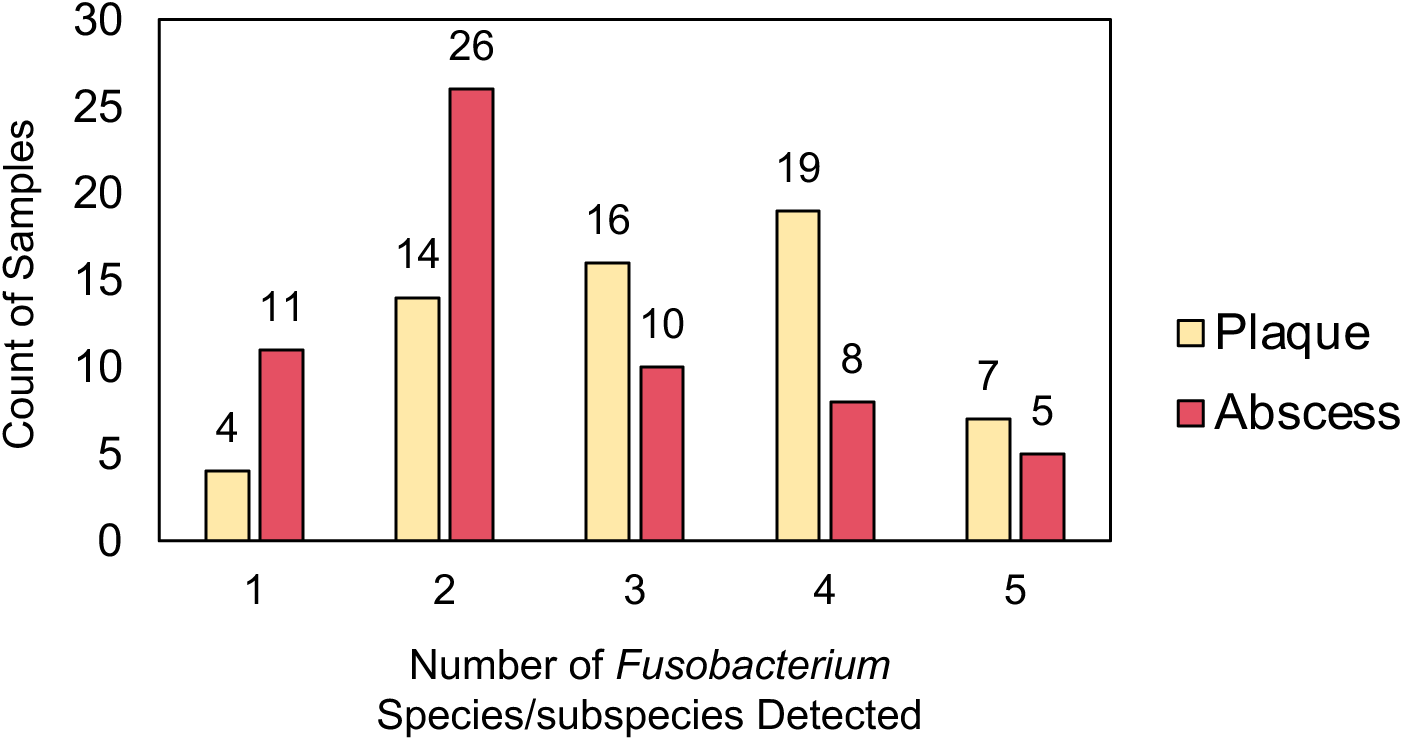
Comparison of oral fusobacterial diversity in plaque and abscess samples. Comparison of fusobacterial species/subspecies diversity in the patient cohort using the NGS identification approach. The detection limit was arbitrarily assigned a threshold of 1% total relative abundance.

The current model of endodontic infections posits that the oral biofilm serves as the ultimate source of infecting bacteria within the tooth pulp chamber, eventually leading to pulpitis and abscessed teeth (22). Accordingly, it would be predicted that the presence of particular fusobacteria within plaque would also correlate with their presence in abscesses (i.e., seed the infections). Thus, the NGS data were further analyzed to determine whether pairwise correlations exist among fusobacteria in the patient specimens. Overall, this prediction was largely supported by the data, as most organisms exhibited positive correlations between their presence in plaque vs. abscess, albeit at varying levels of significance (**Figure 5**). The one major exception was *Fn. polymorphum*, which exhibited a weak negative correlation, presumably due to its substantially lower abundance in abscesses relative to plaque. Conversely, the presence of *Fn*. *animalis* within dental plaque yielded a highly significant positive correlate for its presence in abscesses. A similar positive correlation was noted for *Fn*. *nucleatum* as well. Not surprisingly, both organisms exhibited substantial enrichments within abscesses. We also noted several additional significant correlations between different fusobacteria. For example, the presence of *Fn. polymorphum* in dental plaque yielded a highly significant positive correlation to the levels of *Fn. nucleatum* in the abscess. In dental plaque, *F. periodonticum* is strongly inversely correlated with *Fn. animalis* and to a slightly lesser extent *Fn. vincentii*. In the abscess, *Fn. animalis* exhibits negative correlations with all other fusobacteria, presumably as a consequence of its predominance at this site.

**Figure 5.**
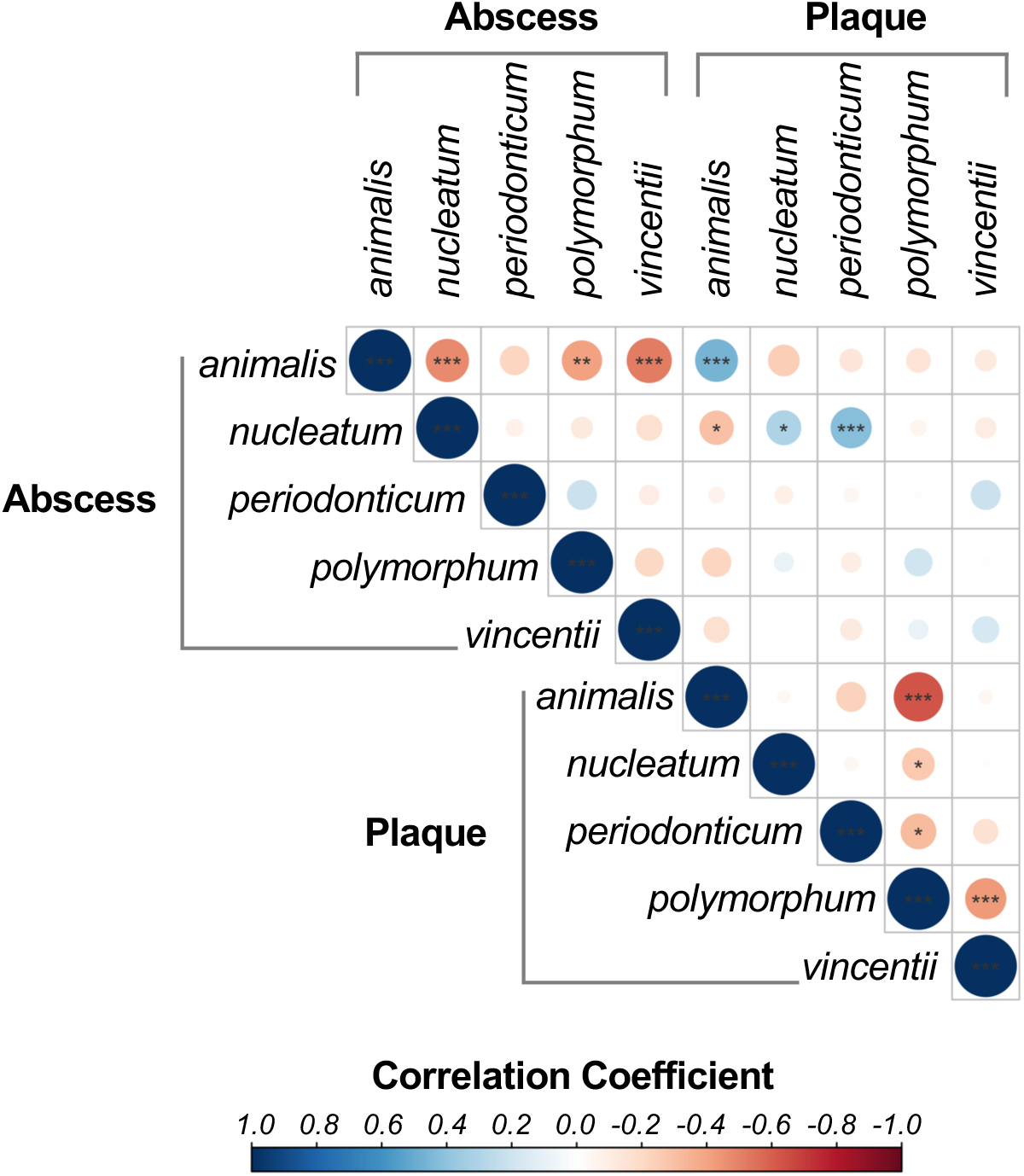
Correlation matrix of species/subspecies co-occurrence in dental plaque and odontogenic abscesses. Blue circles indicate a positive correlation between species in different sites, while red circles indicate a negative correlation. Circles are drawn proportionally to the magnitude of correlation. Significance levels from the Spearman rank correlation test are displayed as asterisks (*p<0.05, **p<0.01, p<0.001). Paired plaque and abscess data from 60 patients were used for analysis.

### 5. Culture-independent qPCR analysis of fusobacterial abundance

Since the aforementioned NGS compositional data were derived from a newly developed methodology, we were interested to verify the validity of these results using qPCR as an independent quantitative approach. We first compared the available *F. periodonticum* and *F. nucleatum* genome data available in NCBI to design a set of universal species-/subspecies-specific qPCR primers (**Table 1).** The specificity of the candidate primers was validated *in silico* using BLASTN alignments (**Table S5**) as well as experimentally by comparing qPCR results of gDNA isolated from representative strains of each type of *Fusobacterium*. As expected, only gDNA from the targeted organisms was robustly amplified by their respective qPCR primers (**Table S6**). Due to the exceptionally low gDNA yields contributed by any single organism within our clinical specimens, we developed a minimal preamplification scheme that facilitated subsequent qPCR analysis for each species-/subspecies-specific primer pair. This approach was validated by generating standard curves of extremely dilute fusobacterial gDNA samples having similar abundances to those found within our clinical specimens. As shown in **Table S7**, the standard curves all yielded correlation coefficients of R^2^>0.98 when using initial gDNA concentrations <0.5 ng µl^-1^ and then further diluting up to 3 orders of magnitude. Encouraged by these results, we next employed the same protocol using a subset of 23 patient-matched clinical specimens randomly selected from the specimen library previously employed for the NGS analyses. The results from these qPCR analyses yielded strikingly similar trends as those previously observed via NGS. *Fn. animalis* exhibited a clear preference for the abscess environment, while *Fn. polymorphum* predominated in the plaque specimens (**Figure 6**). Due to the exceptionally low quantity of gDNA present in many of our clinical specimens, the majority of the *Fn. vincentii*, *Fn. nucleatum*, and *F. periodonticum* samples fell slightly outside of the ranges of their respective standard curves. Nevertheless, when using the standard curves to extrapolate out to their respective qPCR C_T_ values, we observed highly similar proportions as previously determined via NGS analysis (**Figure S2** and **3B)**. In further agreement with the NGS results, qPCR measurements indicated that *Fn. polymorphum* and *Fn. animalis* each contributed >50% of the total fusobacterial gDNA within the plaque and abscess specimens, respectively (**Figure S2**).

**Figure 6.**
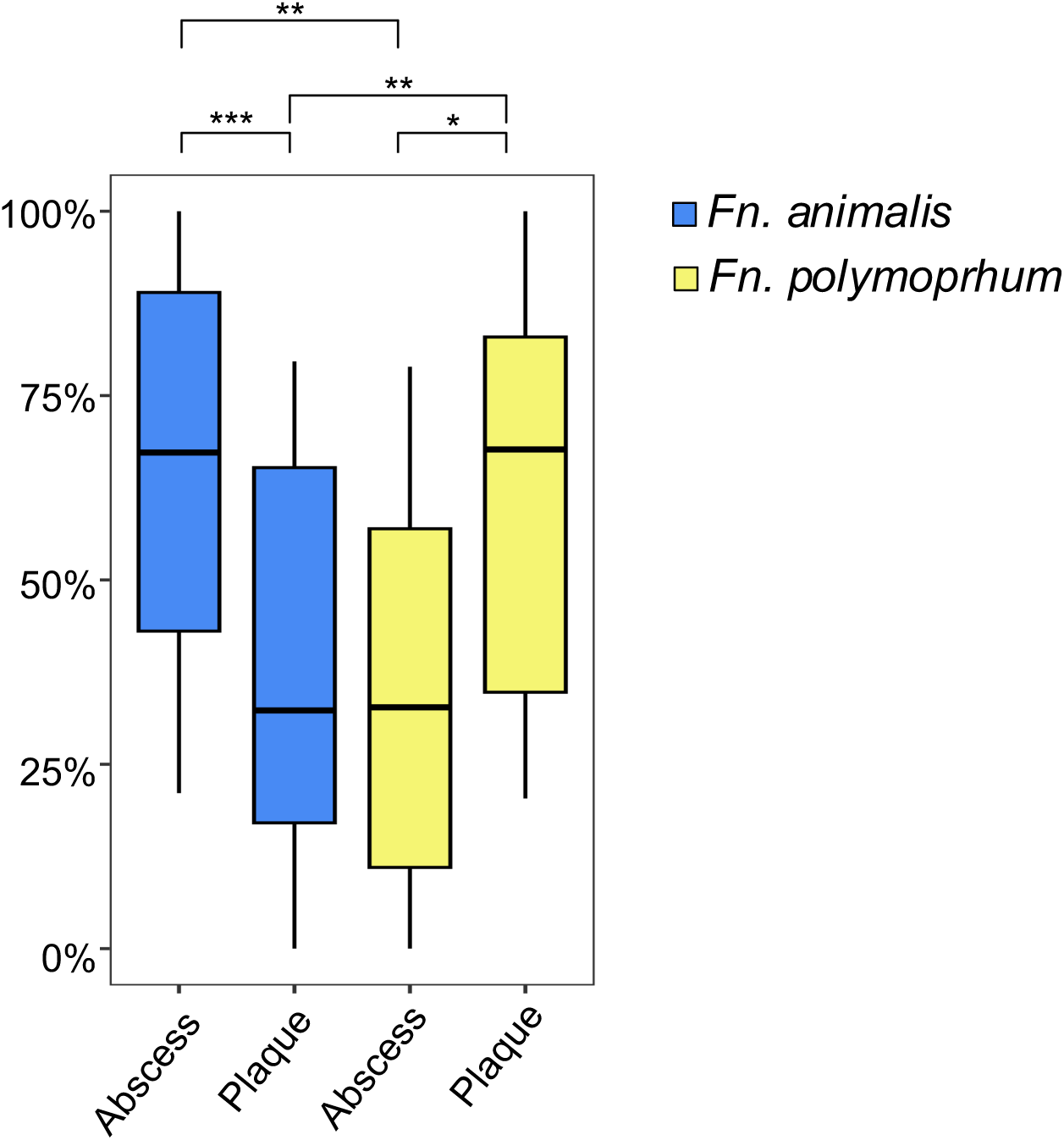
Abundance of *Fn. animalis* and *Fn. polymorphum* in oral plaque and abscess specimens determined by qPCR. A standard curve was used to determine the concentration (ng µl^-1^) of fusobacterial gDNA in each sample. Percentages were calculated based upon the specific concentration of *Fn. animalis* and *Fn. polymorphum* gDNA present in 23 samples and analyzed using pairwise 2-sided t-test comparisons with Bonferroni correction. Significance was determined with a cutoff of 0.05 using Bonferroni adjusted *p*-values (*p<0.05, **p<0.01, ***p<0.001).

### 6. Functional enrichment analysis of *F. nucleatum* and *F. periodonticum* genes

The results of both quantitative genotyping approaches similarly indicated highly biased distributions of oral fusobacteria within sites of health and disease, suggesting that niche-specific ecological preferences exist among these organisms, especially *Fn. polymorphum* and *Fn*. *animalis.* Given their apparent phenotypic differences, we were curious whether this would be reflected in their genomic content as well. Using the Anvi’o pipeline (23), we conducted a functional enrichment analysis to detect Clusters of Orthologous Gene (COGs) groups unique to fusobacterial species/subspecies. Interestingly, genes encoding the high affinity iron transporters FeoA and FeoB were found to be nearly unique characteristics of *Fn. animalis* genomes, with both genes comprising part of the core *Fn. animalis* genome as well (**Table 4**). *Fn. polymorphum* is the only other oral *Fusobacterium* where *feoAB* genes are detectable, but they are only present in <16% of *Fn. polymorphum* strains. It is worth noting that *Fn. animalis* also encodes the same core group of eight iron transport/utilization genes and five iron-associated gene groups that are found in other oral fusobacteria (**Table S8**). In addition, most *Fn. animalis* genomes encode two unique type-1 CRISPR genes, one of which (Cas8b1) is not found in any other *Fusobacterium* genomes surveyed (**Table 4, Table S9**). Interestingly, 11 of the top 20 highest scoring enriched genes were specifically encoded by both *F. periodonticum* and *Fn. polymorphum* (**Table S10)**, which similarly exhibited preferences for growth in dental plaque (**Figure 3**). These genes are primarily associated with central metabolic processes, especially amino acid biosynthesis. As a follow up analysis, we employed the Anvi’o pipeline to compare the average metabolic pathway completeness for numerous biochemical pathways encoded by oral fusobacteria. As expected, *F. periodonticum* and *Fn. polymorphum* uniquely encode complete biosynthetic pathways for multiple amino acids (**Table 5, Table S11**). However, when examined globally, both organisms also yielded the fewest number of missing metabolic pathways overall, suggesting that *F. periodonticum* and *Fn. polymorphum* likely require fewer essential nutrients for growth compared to other oral fusobacteria.

**Table 4.** COG20 gene clusters enriched in *Fn. animalis*.

| COG Function | Functional Enrichment | Adj. q-value | COG Accession | Proportion of Genomes Containing Gene Cluster |  |  |  |  |
| --- | --- | --- | --- | --- | --- | --- | --- | --- |
|  |  |  |  | *Fn. nuc* | *F. per* | *Fn. vin* | *Fn. an* | *Fn. pol* |
| Fe2+ transporter FeoB (FeoB) | 45.8 | 6.43E-08 | COG0370 | 0 | 0 | 0 | 1 | 0.1579 |
| Fe2+ transport protein FeoA | 45.8 | 6.43E-08 | COG1918 | 0 | 0 | 0 | 1 | 0.1579 |
| CRISPR-Cas system type I | 30.6 | 3.70E-05 | COG3649 | 0 | 0.125 | 0 | 0.7857 | 0.1053 |
| CRISPR-Cas system type I-B | 30.6 | 3.70E-05 | COG5938 | 0 | 0.125 | 0 | 0.7857 | 0.1053 |

**Table 5.** Complete metabolic pathways uniquely encoded in *F. periodonticum* and *Fn. polymorphum* genomes.

| DESCRIPTION | PATHWAY | AVERAGE PATHWAY COMPLETENESS |  |  |  |  |
| --- | --- | --- | --- | --- | --- | --- |
|  |  | *Fn. an.* | *Fn. nuc.* | *Fn. per.* | *Fn. pol.* | *Fn. vin.* |
| Cysteine biosynthesis, methionine => cysteine | Cysteine and methionine metabolism | 0% | 0% | 100% | 100% | 0% |
| Cysteine biosynthesis, serine => cysteine | Branched-chain amino acid metabolism | 0% | 0% | 100% | 100% | 0% |
| Ectoine biosynthesis, aspartate => ectoine | Cysteine and methionine metabolism | 0% | 0% | 100% | 100% | 0% |
| Ethylene biosynthesis, methionine => ethylene | Serine and threonine metabolism | 0% | 0% | 100% | 100% | 0% |
| GABA (gamma-Aminobutyrate) shunt | Branched-chain amino acid metabolism | 0% | 0% | 100% | 100% | 0% |
| Summary of total number of missing biosynthetic pathways |  | 9 | 14 | 5 | 3 | 11 |

### 7. Comparative pangenomic analyses of fusobacteria indicate species-level distinctions among *F. nucleatum* subspecies

Due to the unexpected phenotypic and genotypic diversity among *F. nucleatum* subspecies, we were curious to reexamine their taxonomy. We conducted a pangenome analysis using 72 *Fusobacterium* genomes from 7 species/subspecies (**Table S1**), resulting in a total of 6,846 pangene clusters. A phylogenetic tree was constructed using 275 functionally divergent core genes present in all 72 genomes (functional homogeneity index <.95 and geometric homogeneity index of 1) and rooted with *F. periodonticum*/*F. pseudoperiodonticum*. The resulting phylogenetic tree clearly groups the different *F. nucleatum* subspecies into discernable clades that are parallel to other *Fusobacterium* species (**Figure 7**), confirming their genetic relatedness. However, *F. canifelinum*, a mostly non-human species of *Fusobacterium* (24, 25), clusters adjacent to *Fn. polymorphum*, whereas both *Fn. vincentii* and *Fn. animalis* are more distantly related (**Figure 7**), suggesting they may be distinct *Fusobacterium* species. As an additional line of evidence, we calculated the average nucleotide identity (ANI) between the genomes of *F. periodonticum* and the *F. nucleatum* subspecies. An average ANI value <95-96% serves as the standard benchmark for species-level distinctions (26). The ANI within each oral fusobacterial species/subspecies clade was found to be 96%, which is consistent with the clade assignments shown in **Figure 7**. However, when comparing between different oral fusobacteria, the interspecies/subspecies ANI ranged from 85-92% (**Table 6 and Fig. 8**), which is comfortably below the 95-96% threshold used for taxonomic speciation. When considering the weight of independent lines of both phenotypic and genotypic evidence, we conclude that the current *F. nucleatum* subspecies assignments actually represent unique *Fusobacterium* species.

**Figure 7.**
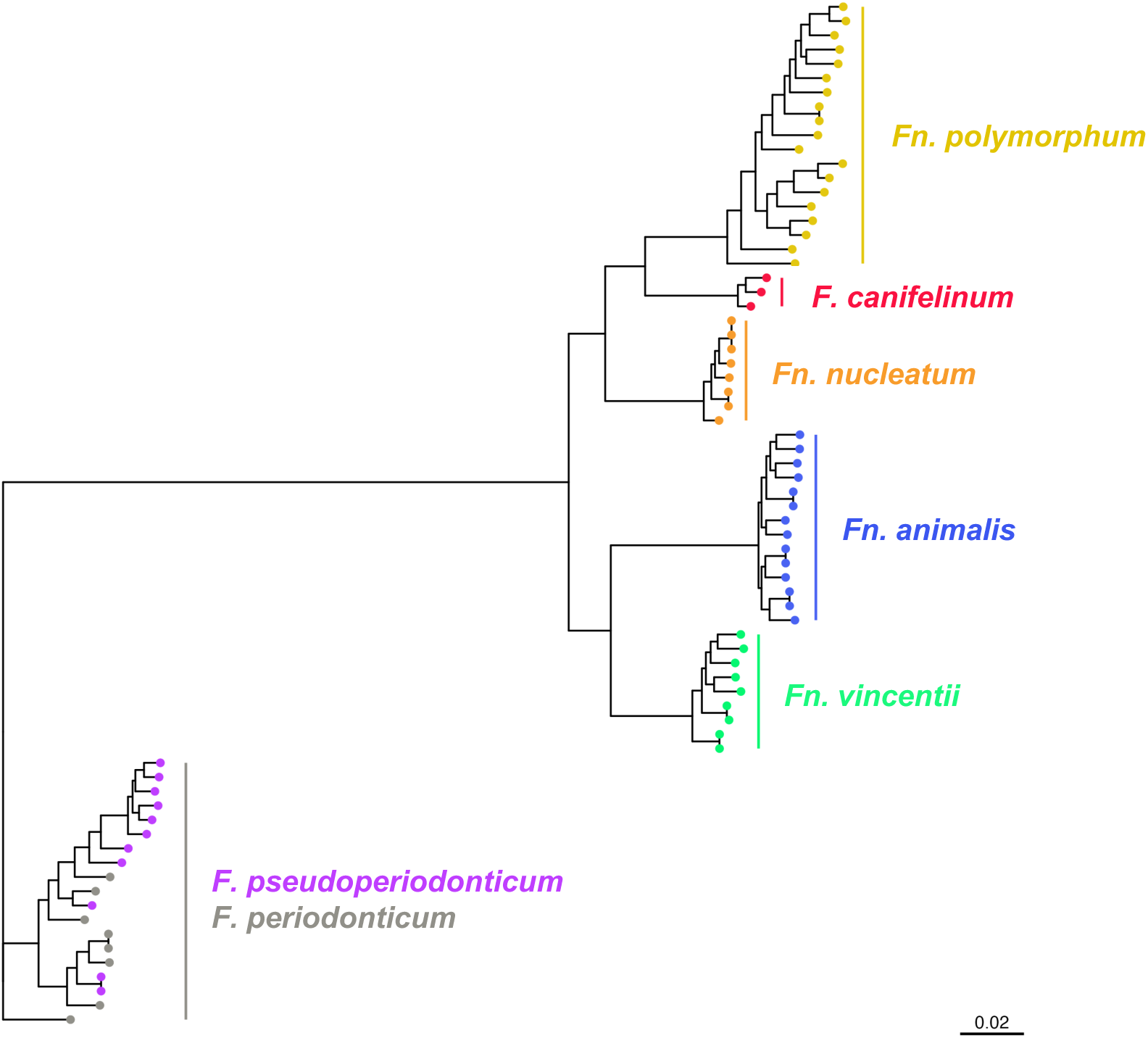
Phylogenetic relationship between *Fusobacterium* species/subspecies. A phylogenetic tree constructed on single-copy core genes having a 0.95 functional homogenicity index was generated in Anvi’o using FastTree. The tree was rooted in the *F. periodonticum/F. pseudoperiodonticum* clade and visualized using ggtree.

**Figure 8.**
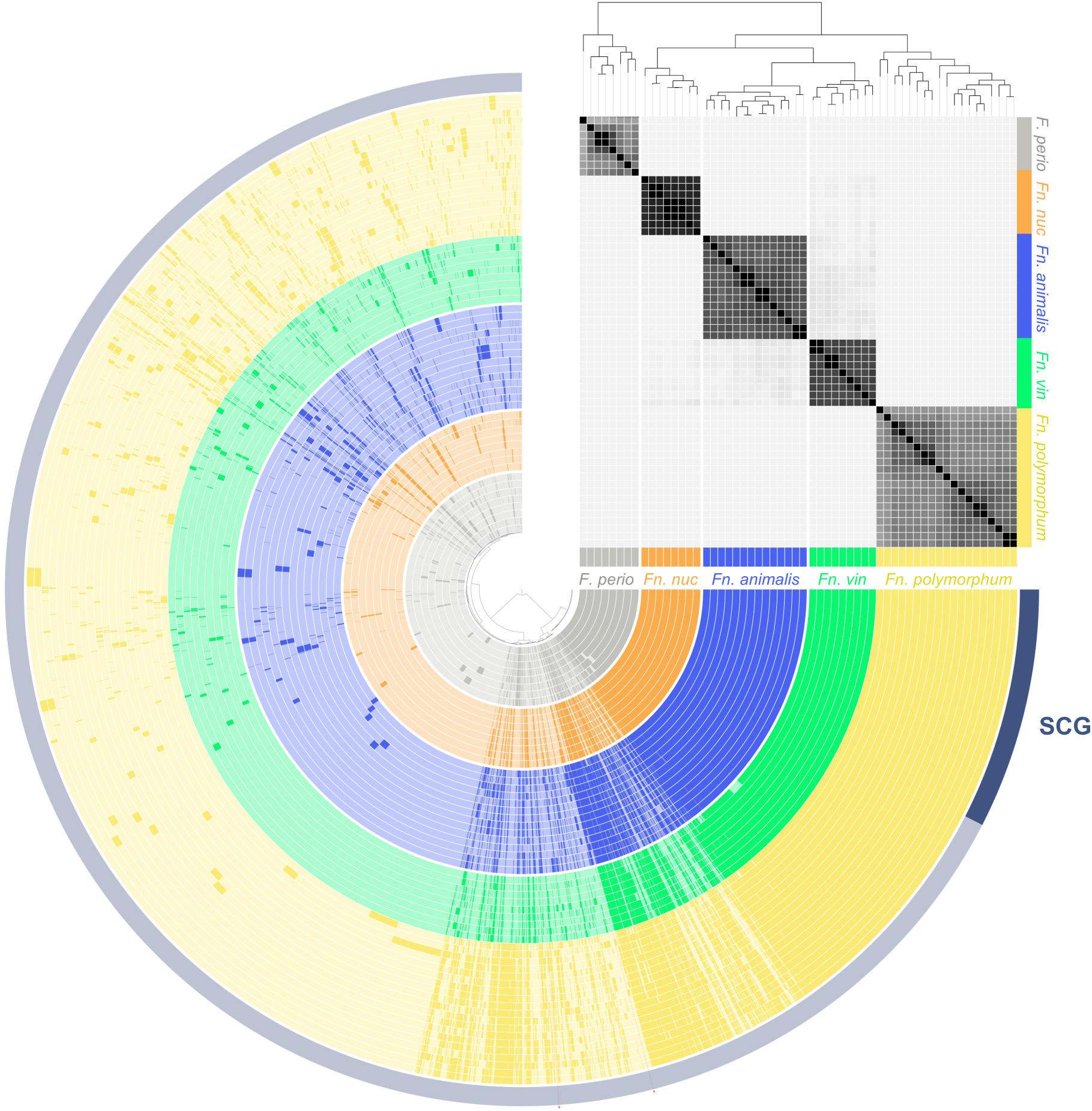
Summary of oral fusobacterial pangenomes, average nucleotide identity (ANI), and phylogeny. The central dendrogram groups gene clusters that are shown around the subsequent rings. Each circular layer represents a unique genome that is colored based upon its subspecies/species annotation. Darker colors indicate the presence of a gene cluster, while lighter shades show the absence of the gene cluster within that genome. A phylogenetic tree, displayed on the top right, was constructed from 233 single copy core gene clusters (SCG) with .95 function homogenicity, a genometric homogenicity of 1, and rooted in the *F. periodonticum*/*F. pseudoperiodonticum* clade. An ANI percentage matrix is displayed below the tree to compare pairwise genome compositions across all genomes. The matrix represents increasing similarity between genomes in progressively darker shades (for ease of visualization, the cutoff for shading was set at 92%). Single copy gene (SCG) gene clusters common to all 58 genomes are shown in dark blue on the outer ring.

**Table 6.** Average nucleotide identity (ANI) between *Fusobacterium* species/subspecies genomes. ANI between genomes from the same species/subspecies are highlighted in yellow.

|  | *Fn. animalis* | *Fn. nucleatum* | *F. periodonticum* | *Fn. polymorphum* | *Fn. vincentii* |
| --- | --- | --- | --- | --- | --- |
| *Fn. animalis* | 97% | 91% | 85% | 90% | 92% |
| *Fn. nucleatum* |  | 99% | 85% | 92% | 92% |
| *F. periodonticum* |  |  | 96% | 86% | 85% |
| *Fn. polymorphum* |  |  |  | 96% | 91% |
| *Fn. vincentii* |  |  |  |  | 98% |

## Discussion

The current study provides an unambiguous assessment of disease association among oral fusobacteria, aided by the unique anatomic aspects of odontogenic abscesses, which are physically sequestered from most other oral bacteria and are only revealed following tooth extraction, assuming the abscesses have yet to rupture and drain. This separation greatly minimizes the potential impact of cross contamination between the patient-matched clinical specimens. To investigate the composition of these complex specimens, we developed three separate approaches useful for detecting and discriminating amongst *F. nucleatum* subspecies. The first employs a qualitative culture-dependent PCR-based approach, while the second uses a quantitative culture-independent NGS approach, and the third employs quantitative subspecies-specific qPCR. Each method has distinct advantages depending upon the desired application. Additionally, this study presents the first subspecies-level assessment of *F. nucleatum* composition within the oral cavity. Both culture-dependent and -independent approaches found *Fn*. *polymorphum* to be the predominant organism present within oral biofilms formed at disease-free sites, whereas odontogenic abscess specimens taken from the same oral cavities exhibit substantial enrichment of *Fn*. *animalis*. Even though *Fn*. *nucleatum* is the *F. nucleatum* subspecies utilized in the vast majority of laboratory investigations, *Fn*. *nucleatum* was the least prevalent subspecies detected in our entire specimen collection, regardless of identification method. Conversely, *Fn. animalis* is largely uncharacterized and has yet to be employed in genetic studies, but our results indicate that this organism strongly predominates at sites of oral inflammation. Furthermore, emerging evidence suggests that *Fn. animalis* likely predominates at extraoral sites of inflammation as well, especially within the colorectal cancer tumor microenvironment (4, 19). Thus, multiple lines of evidence point to *Fn. animalis* specifically as the key fusobacterial organism associated with sites of active inflammation. Accordingly, a variety of compelling evidence indicates that this understudied organism is the most clinically relevant choice for fusobacterial pathogenesis studies of oral and extraoral diseases and should be a major focus of fusobacterial pathobiology research.

Given the distinct health and disease associations of oral fusobacteria, reliably distinguishing amongst them is critical for many applications. Unfortunately, due to the limited number of 16S gene polymorphisms between *F. nucleatum* subspecies (**Figure S1, Table 2**), the commonly employed 16S sequencing approaches only offer relatively low confidence subspecies assignments. To address this limitation, we developed three novel and facile approaches that can be employed for a variety of study designs. For quantitative studies of large clinical specimen collections, the *znpA*-targeting culture-independent NGS approach is clearly preferred and is easily adaptable from typical V3V4 16S workflows. For the quantitation of smaller specimen collections, our qPCR analysis pipeline can yield highly quantitative data from minute quantities of clinical specimen DNA, thus avoiding the extended wait times and high costs typically associated with NGS. For studies requiring *F. nucleatum* strain isolations from clinical specimens, our culture-dependent PCR-screening approach offers a rapid and highly reliable strategy to identify the isolates without having to rely upon sequencing reactions (**Figure 1**). Isolates can be distinguished simply by the presence or absence of particular PCR amplicons using a unique set of PCR primers (**Table 1**) that also function for colony PCR, abrogating the requirement for gDNA extraction. It is worth noting that the primers used for the culture-dependent PCR approach have been verified to function as a multiplex cocktail (**Figure 1**), which further simplifies the identification of strains isolated in pure culture. However, the multiplex PCR approach is not recommended when examining complex mixed samples, as lower abundance isolates can be difficult to detect due to PCR biases favoring the dominant organism(s). In such cases, the primer sets perform far more reliably when used individually. Furthermore, the qualitative culture-dependent approach can be quite useful as a rapid diagnostic of fusobacterial diversity within a complex clinical specimen.

Due to the significant biases of *F. nucleatum* subspecies found at sites of health (plaque) vs. disease (abscess), we were curious whether these apparent phenotypic distinctions would be reflected in their pangenomes and phylogeny. In *Fn. animalis*, we identified several gene clusters that are highly characteristic of this organism, the most prominent of which encodes the high affinity ferrous iron transport system FeoA/B. Orthologs of the FeoAB transporter are crucial for virulence in a variety of organisms, including the oral pathobiont *P. gingivalis* (27, 28) as well as other heavily studied animal and plant pathogens like *Salmonella enterica*, *Helicobacter pylori,* and *Xanthomonas* species (29–32). It is conceivable that in addition to the core collection of iron-transport genes encoded by oral fusobacteria (**Table S8**), the additional high affinity FeoAB iron transporter could provide a specific growth advantage to *Fn. animalis* within the highly inflammatory anaerobic environments where these organisms predominate. In the hypoxic/anoxic conditions of abscesses (33, 34), inflammatory tissue destruction would likely provide a source of reduced ferrous iron, which is specifically targeted by FeoAB (28, 35). The functional enrichment analyses also revealed a surprisingly high number of gene clusters that are specifically enriched in the genomes of both *F. periodonticum* and *Fn. polymorphum* (**Table S10**). Of the 22 genes uniquely encoded by these two organisms, 11 are directly required for the biosynthesis of a variety of amino acids, including methionine, lysine, threonine, cysteine, isoleucine, valine, and leucine (**Table S10**). Likewise, both organisms uniquely encode the predicted molybdenum transporter ModF. Molybdenum is an important cofactor for a number of enzymes, but is especially important in pathways involved in various aspects of sulfur metabolism (36). In both *F. periodonticum* and *Fn. polymorphum, modF* is located in a chromosomal locus containing numerous amino acid biosynthetic genes, including multiple genes listed in **Table S10**, such as *ilvH, ilvC, leuC,* and *leuD*. Likewise, the biosynthetic genes for the sulfur-containing amino acids methionine and cysteine are also included in **Table S10**. Biosynthesis of these two amino acids is often coordinated with a number of sulfur metabolic pathways (37). Molybdenum-containing enzymes are also utilized to reduce oxidized methionine (i.e., methionine sulfoxide) back to methionine (38). Thus, it seems quite likely that ModF is functionally linked with the unique aspects of amino acid biosynthesis found in both *F. periodonticum* and *Fn. polymorphum*. Conspicuously, *F. periodonticum* and *Fn. polymorphum* were both found to favor growth within the typical oral biofilm environment, especially *Fn. polymorphum* (**Figure 3B**). Oral biofilms generally exhibit a greater microbial biodiversity compared to abscesses (39, 40), and are likely to contain a more dynamic and competitive growth environment as a consequence. It is conceivable that the greater metabolic capacities encoded by both organisms (**Table 5, Table S11**) could partially explain their observed growth advantage in dental plaque relative to other oral fusobacteria.

Given the unique phenotypic and genotypic aspects of the *F. nucleatum* subspecies, we were curious to further examine the evolutionary relationships between them, as their distinctions appeared greater than what might be expected from closely related subspecies. ANI has been proposed as the best alternative for the traditional DNA-DNA hybridization method of species classification, with a cutoff threshold value <95-96% ANI indicating unique species (26). The genomes of all *F. nucleatum* subspecies clearly fell below this 95% threshold (**Table 6**). Furthermore, a phylogenetic tree developed from 72 fusobacterial genomes tightly grouped each of the *F. nucleatum* subspecies together, while also indicating that *Fn. polymorphum* is more closely related to the *Fusobacterium* species *F. canifelinum* than to both *Fn. vincentii* and *Fn. animalis* (**Figure 7**). Interestingly, this tree also exhibited parallels to the phenotypic biases observed in our plaque and abscess specimens: *Fn. animalis*, which specifically predominates in abscesses, branches furthest from *Fn*. *polymorphum*, which conversely predominates in the normal oral biofilm environment (**Figure 7**). The pangenome summary displayed in **Figure 8** clearly illustrates the genotypic diversity of the *Fusobacterium* species/subspecies groups. Despite being currently categorized as subspecies of *Fn*, each *Fn* subspecies exhibits comparable genotypic heterogeneity as *F. periodonticum* (**Figure 8**). It has been suggested that *F. nucleatum* subspecies may in fact represent distinct species (41–43), and our results all strongly support this notion. Therefore, to maintain consistency with the extant literature, we propose employing the current *F. nucleatum* subspecies nomenclature as the preferred species names for these organisms (i.e., *F. animalis, F. nucleatum, F. polymorphum,* and *F. vincentii*). As demonstrated in the current study, these organisms exhibit fundamental phenotypic and genotypic distinctions, and should be distinguished accordingly in clinical and laboratory investigations.

## Figures

**Figure S1.**
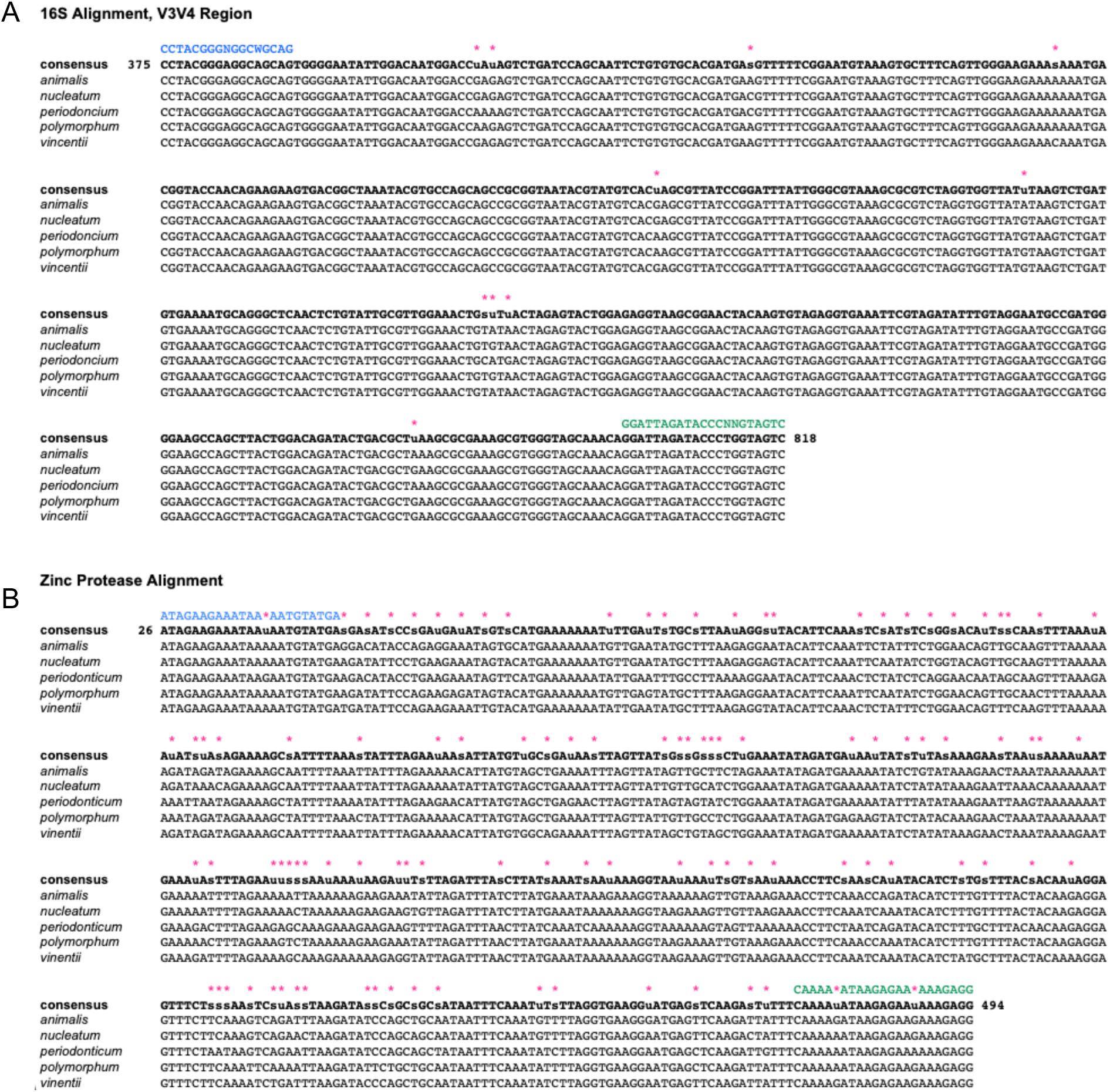
Alignment of the 16S rRNA V3V4 and zinc protease genes (*znpA*) of *F. nucleatum* subspecies and *F. periodonticum.* A consensus sequence is displayed above the species/subspecies alignments. Pink stars above the consensus sequence indicate polymorphisms present in two or more species/subspecies. Primers used to amplify the regions are displayed on the top row in blue and green (reverse-complement). A) The 16S V3V4 region, commonly used for 16S rRNA community analysis, retains almost complete homology between *Fusobacterium* species/subspecies. B) In contrast, the zinc protease (*znpA*) gene sequence displays substantial heterogeneity between species/subspecies.

**Figure S2.**
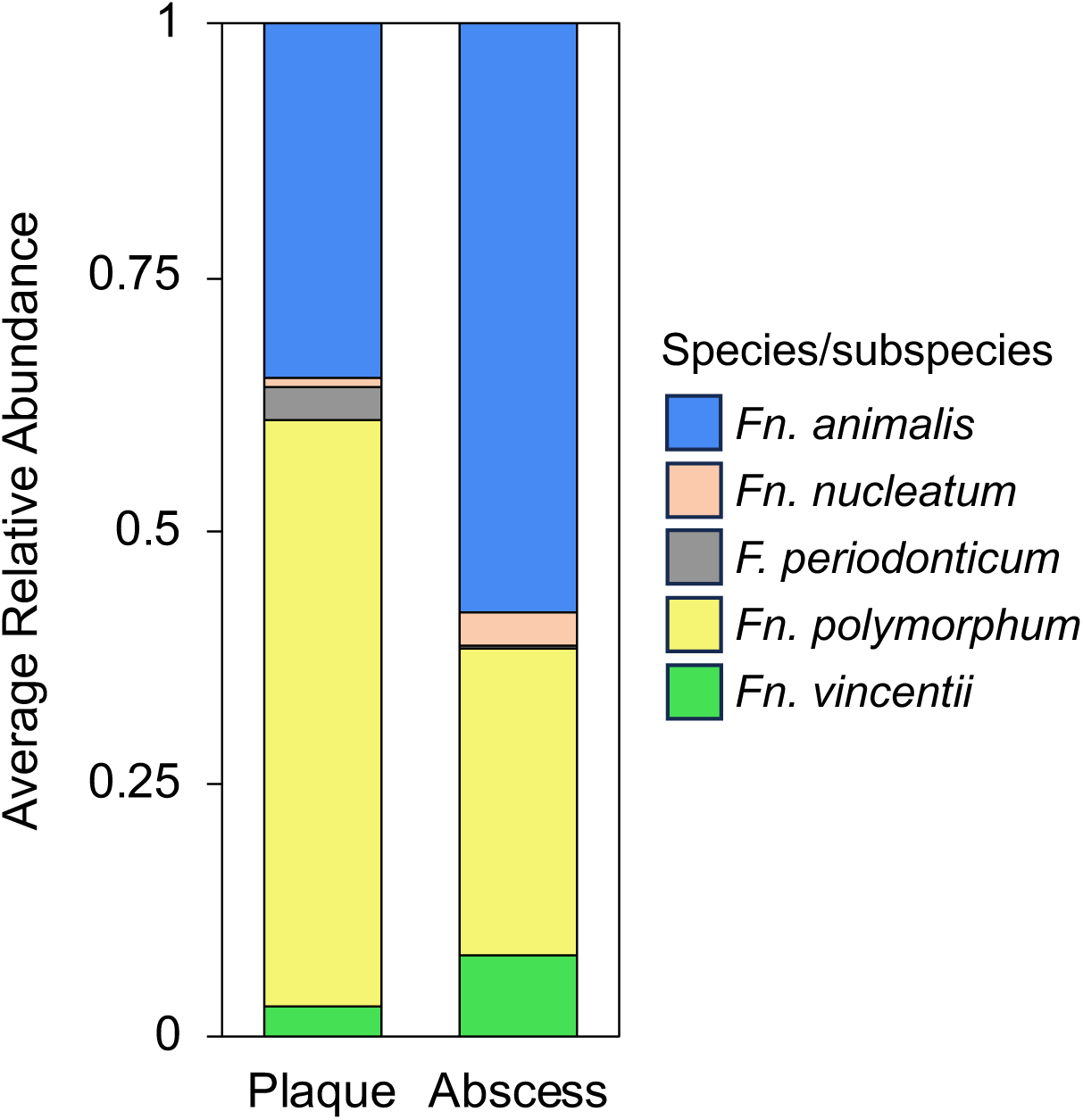
Abundance of *Fusobacterium* species/subspecies in oral plaque and abscess specimens determined by qPCR. A standard curve was used to determine the DNA concentration (ng µl^-1^) of each sample. Proportions were calculated based upon the total concentration of all fusobacterial DNA present from all five species/subspecies in each sample. Results shown are an average of 23 samples.

## Supplementary Table Legends

**Table S1.** Genomes used in this study.

**Table S2.** BLASTN results for species/subspecies-specific primers illustrate *in silico* predicted species/subspecies specificity.

**Table S3.** BLASTN results for zinc protease *(znpA*) outer and inner primer sets indicate their suitability for NGS analyses of oral fusobacteria.

**Table S4.** BLASTN results demonstrate that the zinc protease amplicon can reliably distinguish between oral *Fusobacterium* species/subspecies.

**Table S5.** qPCR primer BLASTN results demonstrate *in silico* predicted oral fusobacterial species/subspecies specificity.

**Table S6**. Validation of species/subspecies sensitivity for qPCR primers.

**Table S7**. Standard curve C_T_ and dilution values for qPCR primers from each species/subspecies.

**Table S8**. Functional enrichment of all iron-associated gene clusters.

**Table S9**. Functional enrichment of CRISPR-associated gene clusters.

**Table S10.** Functional enrichment of gene clusters in *F. periodonticum* and *Fn. Polymorphum* genomes.

**Table S11.** Fusobacterial genomes encoding complete metabolic pathways

## Methods

### Patient sampling and DNA extraction

Deidentified clinical specimens were collected from pediatric dental patients at Oregon Health & Science University dental clinics as part of standard clinical care for one or more abscessed teeth. Prior to the start of the project, all protocols were reviewed by the OHSU IRB and classified as “Not Human Subjects Research”. Briefly, supragingival plaque and extracted tooth abscess samples were collected using cotton swabs and directly placed into sterile, pre-reduced nutrient rich transfer media. In the event of a draining abscess, purulence was sampled from the site of the fistula. Samples were transferred to the anaerobic chamber, mixed with glycerol freezing media, and stored at -80 °C. For the culture-dependent PCR analysis, 100 µl of the freshly collected sample in anaerobic transfer media was spread onto MCDC plates under anaerobic conditions, and incubated at 37 °C for 3-4 days. All growth on the plate was resuspended in sterile saline to an OD_600_ of 1. One ml of that suspension was centrifuged and the pellet was used to extract gDNA using the standard Epicentre DNA extraction kit protocol, except the lysis step was prolonged for 3 hours to ensure total lysis. For NGS analysis, gDNA was isolated from samples using the ZymoBIOMICS DNA Miniprep kit (Zymo Research) per the manufacturer’s instructions, using a double-elution protocol with 50-100 µl of H_2_O warmed to 60 °C without a Zymo-Spin III-HRC filtering step.

### NGS Library Preparation and Sequencing

For the first round of amplification, primers designed to amplify outside the 881 bp region (**Table 1**) were used to generate large initial amplicons. Each PCR reaction contained 0.25 µl Taq Polymerase (Promega), 2.5 µl 10 uM forward and reverse primers, 4 µl 25 mM MgCl_2_, 10 µl 5x Reaction Buffer, 1 µl 10 mM dNTPs, 5 µl DNA, and was adjusted to 50 µl with H_2_O. Amplification was performed using the following thermocycler protocol: 1 cycle of 95 °C for 2 minutes; 28 cycles of 95 °C for 15 seconds, 55 °C for 30 seconds, and 72 °C for 45 seconds; and a final extension at 72 °C for 2 minutes. Amplicons were cleaned with AMPure XP beads (Beckman Coulter) and eluted with 40 µl H_2_O. 20 µl of purified PCR products were used in a second round of PCR to generate the 467 bp zinc protease amplicons with i5/i7 indices for sequencing using the same mastermix protocol outlined above. PCR was performed with the following four-step thermocycler protocol: 1 cycle of 95 °C for 2 minutes; 10 cycles of 95 °C for 15 seconds, 42 °C for 30 seconds, and 72 °C for 45 seconds; 15 cycles of 95 °C for 15 seconds, 65 °C for 30 seconds, and 72 °C for 45 seconds; and a final extension at 72 °C for 2 minutes. Amplicons were cleaned again with AMPure XP beads and indices were attached using the same master mix formulation with the following protocol: 1 cycle of 95 °C for 2 minutes; 8 cycles of 95 °C for 15 seconds, 55 °C for 30 seconds, and 72 °C for 45 seconds; and a final extension at 72 °C for 2 minutes. The resulting DNA was cleaned once again and submitted for MiSeq sequencing at the UC Davis sequencing core (18.3% PhiX spike).

### NGS Analyses

QIIME2 (version 2022.2.0) (44) and DADA2 within the QIIME2 package (45) were used to trim and classify sequences using a feature classifier built from a representative zinc protease gene sequences of each *Fusobacterium* species/subspecies. The classify-consensus-blast command was subsequently used to assign reads to *Fusobacterium* species/subspecies (--p-maxaccepts 1, --p-perc-identity 0.7). R studio Phyloseq (version 1.38.0) (46) was used for subsequent analysis. All samples with fewer than 10,000 reads classified to *Fusobacterium* species/subspecies were excluded from analysis in Phyloseq. Covariance analysis of subspecies/species correlation between plaque and abscess sites was performed using the corrplot (version 0.92) package in R with a Spearman correlation analysis. Ggplot2 in R was used to produced figures.

### Statistical analysis of NGS data

A non-parametric bootstrapping test with 10,000 iterations was used to test differences in proportions of fusobacterial species within plaque specimens, within abscess, or between plaque and abscess. The *p*-values were corrected by Bonferroni correction for the multiple test correction. All computations were performed in R statistical language (47).

### Multiple sequence alignment

16S rRNA V3V4 and zinc protease (*znpA*) sequence multiple sequence alignment was completed using Clustal Omega (48) on the EMBL-EBI webserver with default settings. The *znpA* and 16S V3V4 rDNA gene sequences were obtained from the following strains: *F. nucleatum* subsp. *nucleatum* ATCC 25586, *F. nucleatum* subsp. *animalis* ATCC 51191, *F. nucleatum* subsp. *polymorphum* ATCC 10953, and *F. nucleatum* subsp. *fusiforme* strain ATCC 51190. *F. periodonticum* strain ATCC 33693 was used to generate the corresponding *znpA* sequence, while the partial 16S rRNA gene sequence of *F. periodonticum* strain KP-F10 was used for 16S rRNA V3V4 analysis.

### qPCR

qPCR primers were designed to target genomic locations unique to each species/subspecies. Primer specificity and lack of cross-reactivity was first confirmed using a BLASTN search to identify potential amplicons within species/subspecies (**Table S5**), and further confirmed by qPCR assays on 10-fold dilution series of subspecies/species gDNA (**Table S6**).

Due to the inherently low amount of starting material present in the gDNA samples from clinical isolates, a pre-amplification step with 8 cycles of standard Taq PCR was used to generate sufficient DNA to be detected by subsequent qPCR. Each PCR reaction contained 0.25 µl Taq polymerase (Promega), 2.5 µl 10 µM forward and reverse primers, 4 µl 25 mM MgCl_2_, 10 µl 5x reaction buffer, 1 µl 10mM dNTPs, 1 µl template DNA, and H_2_O adjusted to a final volume of 50 µl. Amplification was performed with the following thermocycler protocol: 1 cycle of 95 °C for 2 minutes; 28 cycles of 95 °C for 20 seconds, 55 °C for 30 seconds, and 72 °C for 30 seconds; and a final extension at 72 °C for 5 minutes. Amplicons were cleaned with AMPure XP beads (Beckman Coulter) and eluted with 25 µl H_2_O. Each qPCR reaction was run with 10 µl Power SYBR Green Mastermix (Applied Biosystems), 0.8 µl forward/reverse primer mix (10 µM), 1 µl DNA, and 8.2 µl H_2_O with the following thermocycler protocol: 1 cycle of 95 °C for 10 minutes, and 40 cycles of 95 °C for 15 seconds and 60 °C for 1 minute. Standard curves were constructed using dilutions of gDNA from each subspecies/species (**Table S7**) pre-amplified for 8 cycles in the same manner as the patient samples.

### Pan-genome analysis

A pangenome was constructed using previously described methods in the Anvi’o workflow (23). For phylogenetic analysis, a pangenome based on 72 genomes (**Table S1**) from 7 *Fusobacterium* species/subspecies was used to identify 275 optimal single-copy core genes present in all genomes with a maximum functional homogenicity index of .95. This collection of genes was used to construct a phylogenetic tree using FastTree (49) within the Anvi’o workflow. A smaller pangenome was created using the same methods from 58 genomes (**Table S1**) from members of the four *F. nucleatum* subspecies groups plus *F. periodonticum*. Functional enrichment analysis (50) and ANI calculations were preformed using the pyANI function within the Anvi’o workflow. Metabolic pathway completeness was estimated using the anvi-estimate-metabolism function within Anvi’o.

## Acknowledgements

Funding for this work was provided by NIDCR/NIH awards DE028252 to J.L.M., DE029612 and DE029492 to J.K.

